# Deep-Worm-Tracker: Deep Learning Methods for Accurate Detection and Tracking for Behavioral Studies in *C. elegans*

**DOI:** 10.1101/2022.08.18.504475

**Authors:** Shoubhik Chandan Banerjee, Khursheed Ahmad Khan, Rati Sharma

## Abstract

Accurate detection and tracking of model organisms such as *C. elegans* worms remains a fundamental task in behavioral studies. Traditional Machine Learning (ML) and Computer Vision (CV) methods produce poor detection results and suffer from repeated ID switches during tracking under occlusions and noisy backgrounds. Using Deep Learning (DL) methods, the task of animal tracking from video recordings, like those in camera trap experiments, has become much more viable. The large amount of data generated in ethological studies, makes such models suitable for real world scenarios in the wild. We propose Deep-Worm-Tracker, an end to end DL model, which is a combination of You Only Look Once (YOLOv5) object detection model and Strong Simple Online Real Time Tracking (Strong SORT) tracking backbone that is highly accurate and provides tracking results in real time inference speeds. Present literature has few solutions to track animals under occlusions and even fewer publicly available large scale animal re-ID datasets. Thus, we also provide a worm re-ID dataset to minimize worm ID switches, which, to the best of our knowledge, is first-of-its-kind for *C. elegans*. We are able to track worms at a mean Average Precision (mAP@0.5) *>* 98% within just 9 minutes of training time with inference speeds of 9-15 ms for worm detection and on average 27 ms for worm tracking. Our tracking results show that Deep-Worm-Tracker is well suited for ethological studies involving *C. elegans*.

## I. INTRODUCTION

*C. elegans*, a free-living nematode, is an important model organism for behavioral studies because of the complex behavioral patterns it exhibits. Researchers have long been attempting to understand the attraction or repulsion of these nematodes (roundworms) to external stimuli such as temperature gradients, food, and chemical signals in studies termed ‘taxis’ [1–7]. In particular, chemotaxis assays have gained importance in understanding the underlying neurological circuits and signalling pathways that govern worm movement towards a particular chemical stimulus [8, 9]. These assays are performed on an agar petri plate by placing worms at a particular distance from a chemoattractant molecule and tracking their movement and trajectory as they move towards the molecule [10–12]. Such assays also find use in identifying and targeting biochemical markers for faster disease diagnosis [13, 14]. A well known method for such assays is the quadrant method [13], where worms are placed in the center of the petri plate from where they migrate to different quadrants on the plate, each containing a test or control chemical molecule. In such experiments, manual counting and marking individual worm trajectory becomes a tedious process. Thus, such studies call for the need of accurate identification of individual worm trajectories and distinguish them from those of other worms. Additionally, other locomotory aspects such as worm posture (coils, turns, omega bends) also need to be quantified. Therefore, accurate detection and tracking of multiple *C. elegans* worms becomes a vital task for ethological studies.

Several tracking softwares based on traditional Machine Learning (ML) and image segmentation algorithms have been proposed earlier [15, 16]. These methods extract individual image frames from video recordings and use thresholding techniques to segment the worm from the background. Such methods perform well in uniform backgrounds where there is sufficient contrast between the worm and its environment. However, laboratory conditions are often noisy and usually such segmenting algorithms fail to accurately segment worms. Moreover, the threshold parameter values have to be manually assigned by the user, which also holds true for most of the ML algorithms. These ML algorithms require step-by-step user feedback to optimize parameters and features needed by the model to train upon. Such methods also breakdown upon lack of well defined features: for example worm size can be an ill-defined feature for a model because of phenotypic differences between worms in different developmental stages and can also be caused due to microscope magnification changes. A classic example of this approach was implemented in the Tierpsy Multi-Worm behavior tracker [17, 18] which uses adaptive thresholding to distinguish worms from the background based on light intensity. The filter parameters are user dependent and need to be changed for different experimental settings making it unsuitable for uneven lighting conditions or heterogeneous backgrounds. The software also fails to keep continuous tracks of multiple worms during occlusions and trajectories are reinitialized after the worms separate out. Other ML algorithms like the one implemented in Multi Animal Tracker by Itskovits *et al*. [19] has issues with misidentifying animals, as evidenced in video results on the model organism zebrafish, where the tracker incorrectly detects fish reflections as actual fish instances.

Deep Learning (DL) approaches, on the other hand, make use of the power of Convolutional Neural Networks (CNNs), which can independently extract features of an object while maintaining the object’s spatial configuration. These models are well suited for ethological studies because of the large amount of video and image data generated from behavioral experiments. Although, these models can require longer training times, they outperform traditional methods during test time providing near real-time track results. DL models also require minimum user interference making them versatile even in changed experimental setups. With availability of powerful graphic cards (GPUs) in devices as well as remote access via cloud computing, such models can be easily deployed even in small devices like Raspberry Pi. Present times have seen an increase in the implementation of DL models in tracking animals [20–22]. These models are highly accurate and can keep tracks of multiple unmarked animals in a frame. However, a real-time, accurate and easy to deploy tracker with minimum preprocessing and training time is still required.

Deep-Worm-Tracker is a step forward in this direction and combines two state-of-the-art DL models — You Only Look Once (YOLOv5), an object detection model [23] and Strong Simple Online Real Time Tracking (Strong SORT) as a tracking backbone [24]. The YOLO family of object detection models [23, 25–29] are single step detectors with accuracy of these models at par with that of the Region based Convolutional Networks (R-CNN) family of models [30–32]. YOLO does not employ a Region Proposal Network (RPN) like that of the R-CNN models which makes it faster during inference times, thereby making it suitable for live camera tracking experiments. In the context of animal tracking, the YOLO detection models have previously been used in tracking livestocks [33, 34]. Therefore, we chose YOLOv5 as our object detection model for detecting *C. elegans* worms.

Oftentimes, object detection models go wrong and misidentify worm instances during rapid frame drops or during longer periods of worms occlusions. To account for this, we combine Strong SORT, which is an improvement to the earlier Deep SORT model [35], as a tracking backbone. Strong SORT uses a combination of motion based and appearance based features extracted by a deep network to keep track of multiple worm IDs. A shallow deep network is trained on a worm re-ID dataset to account for occlusions and preserve worm IDs after the worms are no more occluded. This task is exceptionally challenging for worm or animal tracking because animals, unlike humans, do not have distinguishing physical appearances such as varying clothing or hairstyles. Such re-ID datasets are currently scarce and limited across species, as reported in the work by S Schneider *et al*. [36]. The article also illustrates that DL algorithms perform well in distinguishing two fruit flies based on learnt features, that are humanly impossible to discern. In light of this, we curated a worm re-ID dataset containing 32 worm identities, which to the best of our knowledge is first of its kind for *C. elegans*.

We find that the YOLOv5 detection model achieves high mean average precision *>* 98% with minimum training time of 9 minutes with just few annotated video frames. We also show that the Strong SORT model trained over the re-ID dataset acts as a supporting backbone and achieves lower ID switches between multiple worms. The entire pipeline of the Deep-Worm-Tracker along with the training datasets is available at the github repository (https://github.com/knaticat/Deep-Worm-Tracker). Our Deep-Worm-Tracker model can also be expanded for detecting and tracking other model organisms in real-time based on the researcher’s choice.

The rest of the article is organized as follows: In Section II, we discuss the methods used to carry out data collection, dataset preparation, model used and related training and evaluation metrics. In Section III, we present the results/performance of the Deep-Worm-Tracker. Finally, we present a discussion of our results in Section IV.

## II. METHODS

This section discusses the methods used in (A) data collection, (B) image acquisition, dataset preparation, (D) model configuration and setup, (E) evaluation metrics used and (F) training setup.

### *A. C. elegans* strains and culture methods

The *C. elegans* worms used in this study were chosen at different stages of development right from larval stages (L1-L4) to adult forms in order to include developmental diversity in the study. Strains of *C. elegans*: FK101, N2, EG7985 were used for culturing and were maintained at 22^◦^C. The worms were grown on fresh Nematode Growth Media (NGM) agar plates. These plates were seeded with *E. coli* OP50 lawns as a food supply for worms. The seeded bacterial lawn makes worm picking easier and helps keeping the worms in focus throughout image acquisition.

### B. Image Acquisition

For image acquisition, fresh agar plates were used and worms were transferred to these plates. Images were acquired using a Leica S9i stereo microscope. The camera used for the setup was CMOS camera and images were taken at a resolution of 0.8MP, 2MP and 5MP. To make Deep-Worm-Tracker adaptive to different magnifications, images were also collected at varying magnifications of 1.5X, 2X and 5X. The worm re-ID dataset was prepared by gathering three to five 30-second video recordings at 100 FPS for each worm. Cropped images of worms were obtained from these video tracklets using detections from the YOLOv5 model at an interval of 5 frames giving ∼ 60 images for each tracklet.

### C. Dataset Preparation

Deep-Worm-Tracker is trained upon two separate worm datasets. The first, worm detection dataset, is used for training the YOLOv5 object detection model. We extracted individual frames from 30s video recordings and selected frames containing multiple worms (1-15) as positive instances for manual annotation. We employed Roboflow, an online annotation tool for generating annotation files for the YOLOv5 model used in Deep-Worm-Tracker. We feed tiled data augmented images while training our detection model, considering the small pixel dimensions of worms in high resolution video files. In addition to this, frames comprising of empty plate images with worm trails or plates with dust particles were added into the model while training. These negative samples provide feedback to the model and help the model in distinguishing features it should not detect. Next, we provide a worm re-ID dataset containing 32 different worm identities. Each of the worm identities contain ∼ 100 images acquired by cropping detection outputs obtained from the YOLOv5 model. The image files were renamed according to the nomenclature used while preparing the Motion Analysis and Re-identification Set (MARS) dataset [37]. Some sample images in this dataset are shown in Fig. 1

**Figure 1:**
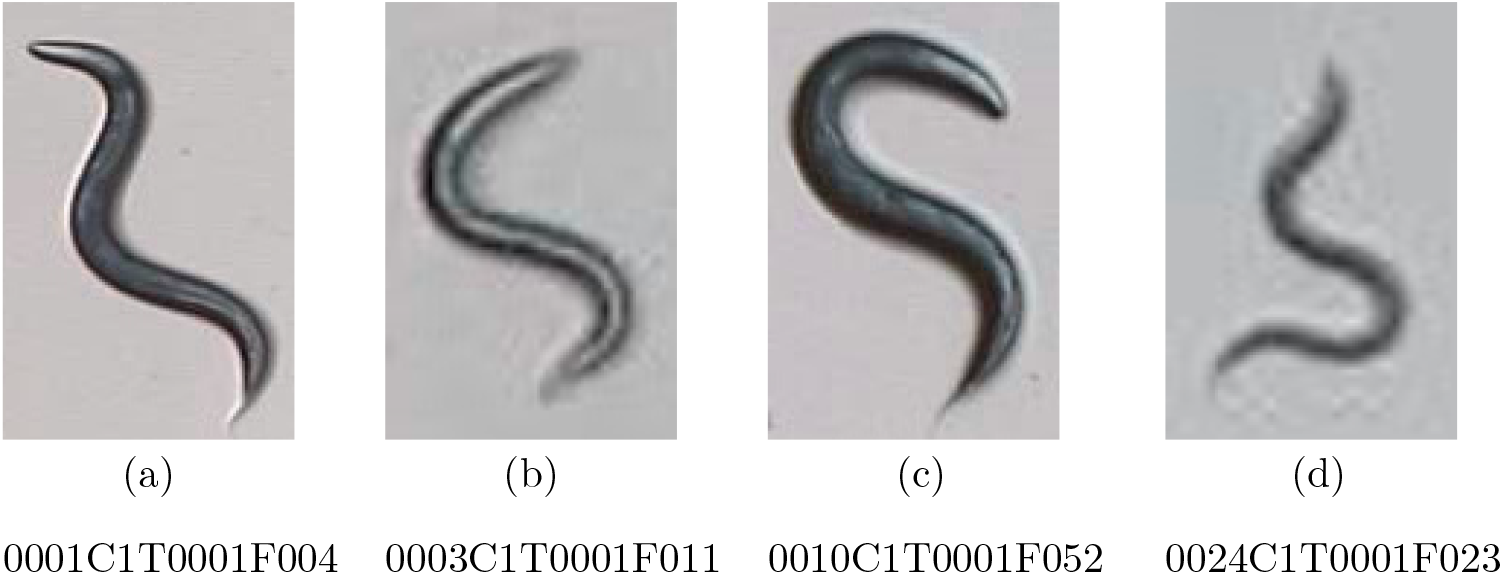
Sample images of different worm identities from the worm re-ID dataset

This worm re-ID dataset is used in training a shallow deep network used in the Strong SORT tracking model. The Strong SORT tracking model uses appearance feature maps of each worm identity extracted using a deep CNN. These appearance features are then combined with the motion features available by the Kalman filter to keep track of multiple worms. A combination of worm appearance and motion features helps address the worm occlusion problem.

### D. Model Configurations

The network dimensions used for training the YOLO detection model was set at values of 640 × 640, 832 × 832 and 1024 × 1024. The best detection results considering a trade off between detection speed and accuracy was observed for the network dimension of 832 × 832. The model was trained for 30 epochs. Pre-trained YOLOv5 weights, trained on the Microsoft Common Objects in Context (MS-COCO) dataset [38] were used during training. The worm detection dataset was split in a 80:10:10 split with 1407 images being used for training, 180 validation images and 171 to be used as a testing set. For training the Strong SORT tracking model, we used the training implementation code as given by torchreid. We trained the Strong SORT model for 100 iterations and monitored it’s performance in tensorboard till the accuracy converged to *>* 95%.

### E. Evaluation Metrics

To evaluate the performance of the YOLOv5 object detection model the following metrics were used: Recall or sensitivity defined as the ratio of all True Positives (object is correctly identified) to the total of True Positives and False Negatives (missed instances of an object) 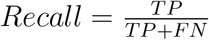. Precision or specificity defined as the ratio of True Positives to the total of True Positives and False Positives (incorrect identification as being an object) 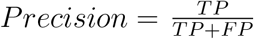. Mean Average Precision (mAP) is the mean of the average precision for each class. Average precision is defined as the area under the precision-recall curve. Usually the mAP@0.50 indicates the mean average precision at an IoU threshold of 50% overlap between the predicted and ground truth bounding boxes. F1 Score is a combination of precision and recall as a single value defined by the harmonic mean of the two values, i.e., 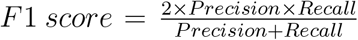. We also use the inference time (ms) or the FPS of the final tracked video to test the speed of the Deep-Worm-Tracker model. The Intersection over Union (IoU) score is used to measure how much of the predicted bounding box overlaps with the actual ground truth box. It is mathematically defined as the ratio of the area of overlap between the two boxes to their combined areas. 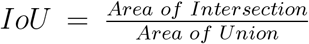 We set the IoU threshold to 0.5 for the final tracked frames. For evaluating performance of the Strong SORT tracking model we used the Multiple Object Tracking Accuracy (MOTA) metric which is defined as: 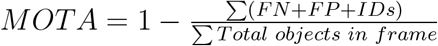

### F. Training Setup

The Deep-Worm-Tracker model was trained on a Linux 20.04 Operating System. The GPU used for training the models was a NVIDIA Quadro RTX 4000 with a 8GB RAM. The models were implemented using the Pytorch machine learning framework (Pytorch 1.12.1). In addition, the models require support from the CUDA (Version 11.4) and CUDNN libraries available from NVIDIA.

## III. RESULTS

More often in laboratory settings, uneven lighting conditions and ink markings left on the petri plate to demacrate different quadrants, account for background noise in chemotactic assays. Added upon this are the worm trails and worm picking marks left on agar plate that can be easily misidentified as an actual worm by traditional ML algorithms. Moreover, when plates are exposed outside during such assays, dust particles from the outer environment often settle upon the agar media increasing the chances of misidentifying actual worms from these false positives. Additionally, an ideal tracker should work using markerless IDs on animals to make the behavioral assay non-invasive. All these factors demonstrate the difficulty in identifying an actual worm instance in an image frame, with this problem raised immensely when multiple worms are used for tracking. While tracking multiple worms, the problem of correctly re-identifying the same worm after a worm occlusion also poses a challenge.

Deep-Worm-Tracker achieves real-time, accurate tracking results under fewer image frame annotations and faster training time. It can be used both for offline as well as online markerless video tracking experiments. Video files, images or live webcam recording can be used as input to the tracker. Deep-Worm-Tracker first runs the YOLOv5 detection model to detect worm instances in each frame. The worm predictions are then passed on to the Strong SORT tracking model that extracts appearance and motion based features from these predicted boxes and using the Hungarian assignment algorithm keeps the identity of each worm preserved across frames. Deep-Worm-Tracker also provides posture data in the form of worm segment and worm skeleton. Segmentation and skeletonization algorithms are often applied to the entire image frame during testing, accounting for a lot of background noise in the outputs by these methods. Instead, Deep-Worm-Tracker uses the predicted bounding boxes from the detection and tracking modules as an input to apply segmentation and skeletonization only in these confined regions. In this manner, Deep-Worm-Tracker greatly reduces the chances of background noise being captured by the model. The representative flowchart of this method is shown in Fig. 2. Deep-Worm-Tracker also outlines the trajectory taken by each tracked worm along with their IDs. This is suitable for studies that focus on the exact movement pattern of *C. elegans* worms. The entire workflow of Deep-Worm-Tracker is illustrated in Fig. 3.

**Figure 2:**
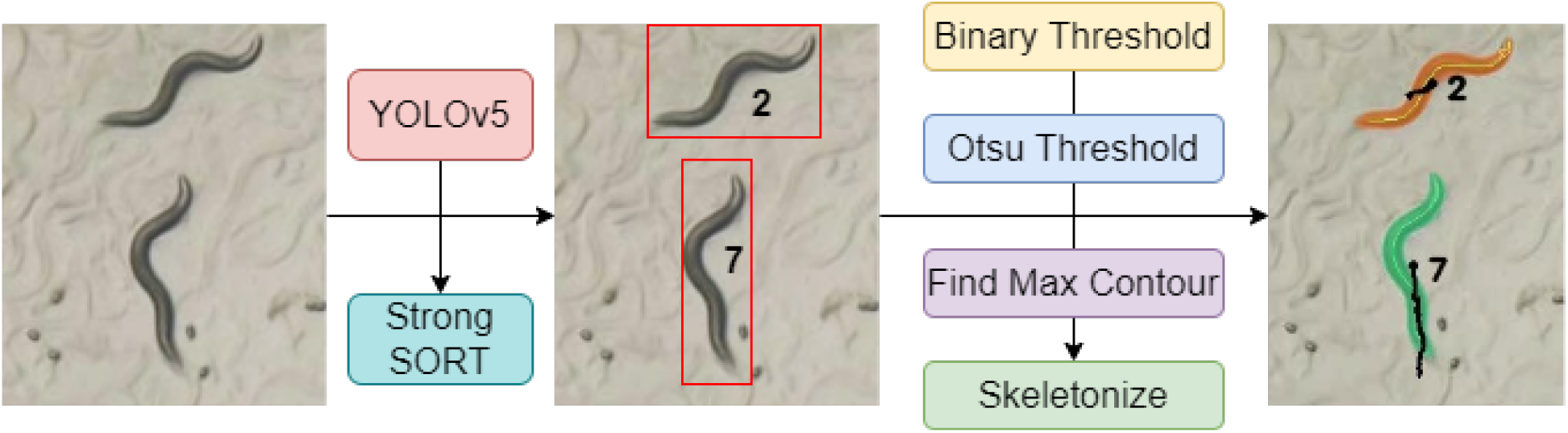
Segmentation and Skeletonization techniques used by Deep-Worm-Tracker. The predicted bounding boxes from the YOLOv5 and Strong SORT models are used as input regions for applying threshold segmentation and skeletonization algorithms. This avoids capturing unwanted background noise.

**Figure 3:**
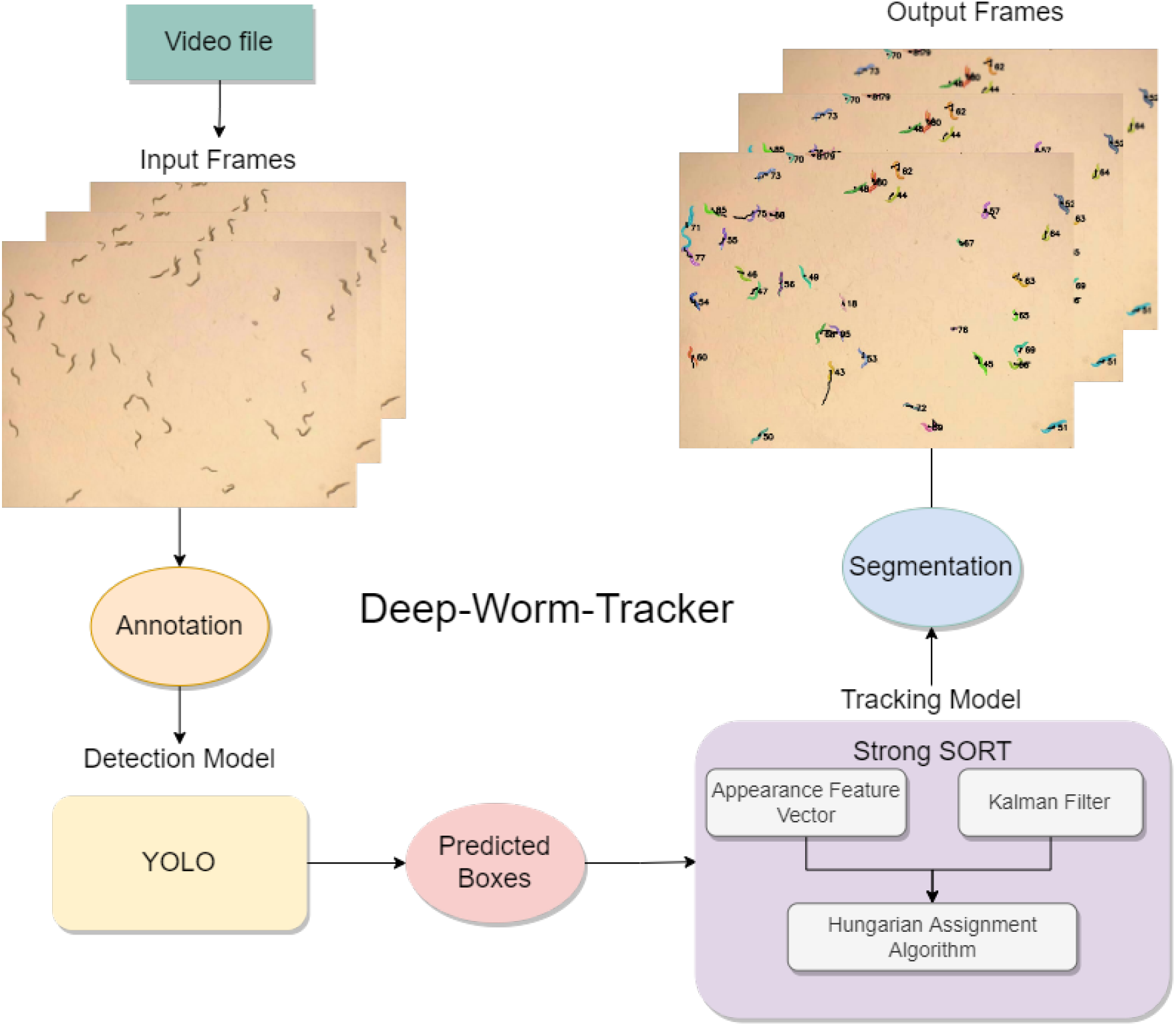
The workflow of Deep-Worm-Tracker: Input video files are first extracted frame by frame from the model. Next, these input image frames are resized according to the network dimension of the YOLOv5 detection model. These resized images are then passed on for detecting worm instances. The predicted bounding boxes of each worm is then used as input to the Strong SORT tracking model. Strong SORT uses appearance features extracted by a shallow re-ID network and combines it with the motion based features from the Kalman Filter to retain individual worm IDs. Upon completion of the tracking steps, the segmentation and skeletonization algorithms are called upon these predicted regions

Based on the worm detection algorithm, Deep-Worm-Tracker achieves a mean aver-for each of the worm IDs to give the final tracked and segmented results. age precision at 0.5 IoU threshold (mAP@0.5) of *>* 98% in the first 15 epochs itself. This mAP@0.5 is achieved at a minimum training time of ∼ 9 minutes for the simplest model. The trained models have an almost equal precision and recall values, illustrating the balanced design of the detection algorithm for our use case. In order to avoid overfitting, we also evaluate the training and validation loss curves that can be found in SI. For testing our model, we chose the best model weights that are automatically stored by the YOLOv5 model while training. We trained YOLOv5 object detection models (nano and small) on varying network image dimensions (640 × 640, 832 × 832, 1024 × 1024). The best results in terms of a trade-off between detection accuracy and inference speed was obtained for a network dimension of 832 × 832. On using these network dimensions, the Deep-Worm-Tracker provides a real-time inference speed of 9-15ms based on the duration of the video and number of worms (5-40 worms for testing). The training curves illustrating the precision, recall, mAP@0.5 and training time are shown in Fig. 4. As evident from the results, although the YOLOv5 detection model is sufficient to give accurate and real-time detection results, a tracking model is still required for cases of missed detections and frame drops. Previous study by K Bates *et al*. [39] used a Faster-RCNN based detection model solely without a tracking backbone for worm tracking. Also, the detection model alone cannot account for long term worm occlusion while tracking multiple worms. Deep-Worm-Tracker combines the YOLOv5 detection model with Strong SORT which correctly re-identifies worm IDs in cases of missed detections during some frames. Deep-Worm-Tracker trained on our manually curated worm re-ID dataset achieves a training accuracy *>* 90% within the first 50 epochs. We trained versions of osnet and mobilenet re-ID models available through the Torchreid package [40]. These lightweight models support deployment of our tracker in devices like Raspberry Pi or CPUs which do not have access to GPUs, while still providing near real-time tracking results.

**Figure 4:**
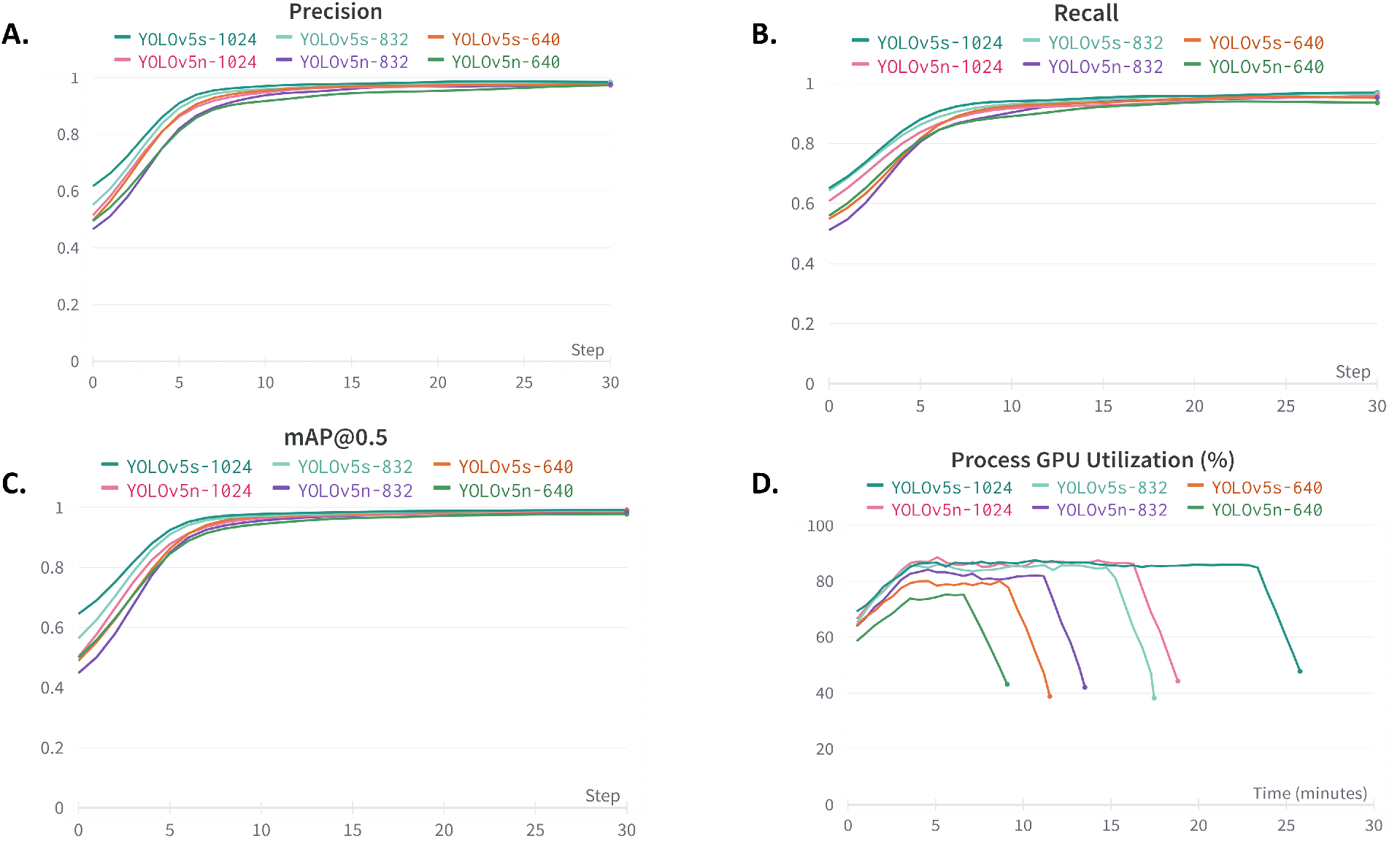
Training Curves for the YOLOv5 worm detection model. (A) The precision curves (B) The recall curves (C) The mAP@0.5 curves (D) The process GPU utilization and training time curves for different input network dimensions for the YOLOv5 nano and small models.

We tested Deep-Worm-Tracker for tracking on four different video recordings of single and multi worms (Video files are provided in SI). The results are presented in Fig. 5 as snapshots of the video recordings used for testing Deep-Worm-Tracker.

**Figure 5:**
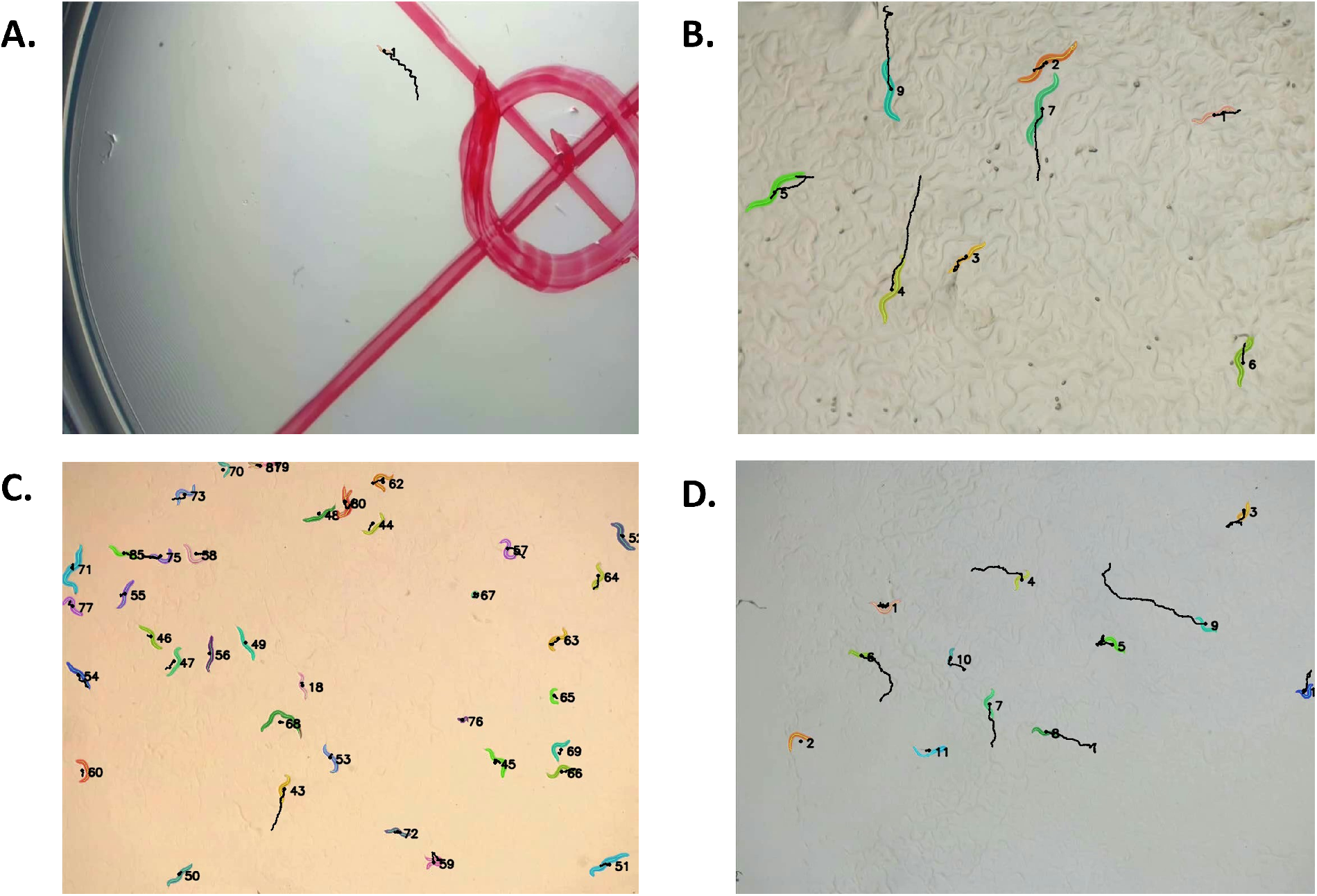
Results of Deep-Worm-Tracker on (A) single and (B-D) multi worm tracking recordings. Individual worm IDs and trajectory marks are also shown along with worm skeleton and segmentation masks.

We also summarize the inference speed of our Deep-Worm-Tracker model on these video recordings in Table I.

**Table I:** Summary statistics of various video files used for evaluating the performance of Deep-Worm-Tracker

| Video File<br>(.mp4) | No. of<br>Individuals | Duration of<br>Video | YOLOv5<br>Inference Speed | Strong SORT<br>Inference Speed | FPS | MOTA |
| --- | --- | --- | --- | --- | --- | --- |
| worm1 | 1 | 30s | 9.3ms | 16.8ms | 29.5 | 99.60% |
| worm2 | 40 | 0-10s | 14.2ms | 37.8ms | 19.2 | 70.44% |
|  | 44 | 10-20s |  |  |  | 63.16% |
| worm3 | 9 | 0-10s | 10.4ms | 26.4ms | 27.1 | 85.66% |
|  | 9-10 | 10-20s |  |  |  | 70.88% |
|  | 9 | 20-30s |  |  |  | 47.70% |
| worm4 | 11 | 0-10s | 11.1ms | 25.2ms | 27.5 | 82.18% |
|  |  | 10-20s |  |  |  | 69.63% |
|  |  | 20-30s |  |  |  | 34.90% |

Based on the MOTA tracking accuracy we find that Deep-Worm-Tracker performs well in keeping track of single worm instances. In cases of multi worm tracking, Deep-Worm-Tracker is able to keep track of multiple unoccluded worms. However, our model finds it difficult to keep track of worms that leave or enter the field of view of the camera or during heavy occlusions which don’t get resolved. In summary, Deep-Worm-Tracker can be deployed for multi worm tracking and find use in chemotaxis assays in *C. elegans*.

## IV. DISCUSSION

In this study, we propose Deep-Worm-Tracker, a tracking model for tracking multiple *C. elegans* worms for chemotactic behavioral studies. Keeping track of multiple worms during chemotactic assays becomes a tedious task if performed manually. Moreover, traditional ML techniques are highly susceptible to noisy background conditions and require a lot of user dependent parameter tuning for getting accurate animal tracks. In light of this, we used two current state-of-the-art DL models, YOLOv5 for worm detection and Strong SORT as a tracking backbone to our Deep-Worm-Tracker. We manually curated a worm detection dataset used for training our YOLOv5 object detection model. In addition, we also present a first-of-its-kind *C. elegans* re-ID dataset consisting of 32 worm IDs used for training the Strong SORT model.

We tested Deep-Worm-Tracker using single and multi worm video recordings. We find that Deep-Worm-Tracker can accurately keep track of multiple worm IDs and incorrect or missed IDs arise when worms are partially visible, as is the case when worms enter the field of view of the camera or when the worms completely overlap with each other. In such cases of extreme overlaps, Deep-Worm-Tracker assigns new IDs to worms after they separate from heavy occlusion in later frames. We suggest that these cases can also be resolved with more training images provided by the user. Deep-Worm-Tracker could also correctly identify actual worm instances from other background noise such as worm trails [video-file:worm2.mp4] or thread like dust particles [video-file:worm4.mp4] left on the agar plate. Deep-Worm-Tracker is also invariant to magnification changes and lighting conditions and is well adapted to be used for behavioral studies in laboratory conditions.

In conclusion, we find that Deep-Worm-Tracker can be useful for ethological studies involving chemotaxis in *C. elegans*. We have also enabled Deep-Worm-Tracker to extract worm skeleton and segmentation data from tracked worms to be used for genetic and neurological assays in *C. elegans*. Our proposed tracker also supports visualization of individual worm trajectories making it easier to study *C. elegans* locomotion. Deep-Worm-Tracker requires minimum installations and can be easily run on a cloud computing platform like Google Colab with access to GPU. Finally, our study is not limited to a single model organism in *C. elegans* and can be extended to incorporate other organisms based on a researcher’s choice for use in other behavioral studies.

## DATA AVAILABILITY

The data that supports the findings of this study are available within the article and its Supplementary information files. The entire pipeline of the Deep-Worm-Tracker is available at the github repository https://github.com/knaticat/Deep-Worm-Tracker.

## ACKNOWLEDGMENTS

This work is supported by the Science and Engineering Research Board (SERB) POWER Research Grant (Ref. No. SPG/2021/002732) awarded by the Department of Science and Technology (DST), India.

