## Supplementary Information for "Deep-Worm-Tracker: Deep Learning Methods for Accurate Detection and Tracking for Behavioral Studies in *C. elegans*"

1. Additional training Curves for the YOLOv5 worm detection model. The input network dimension is $832 \times 832$.

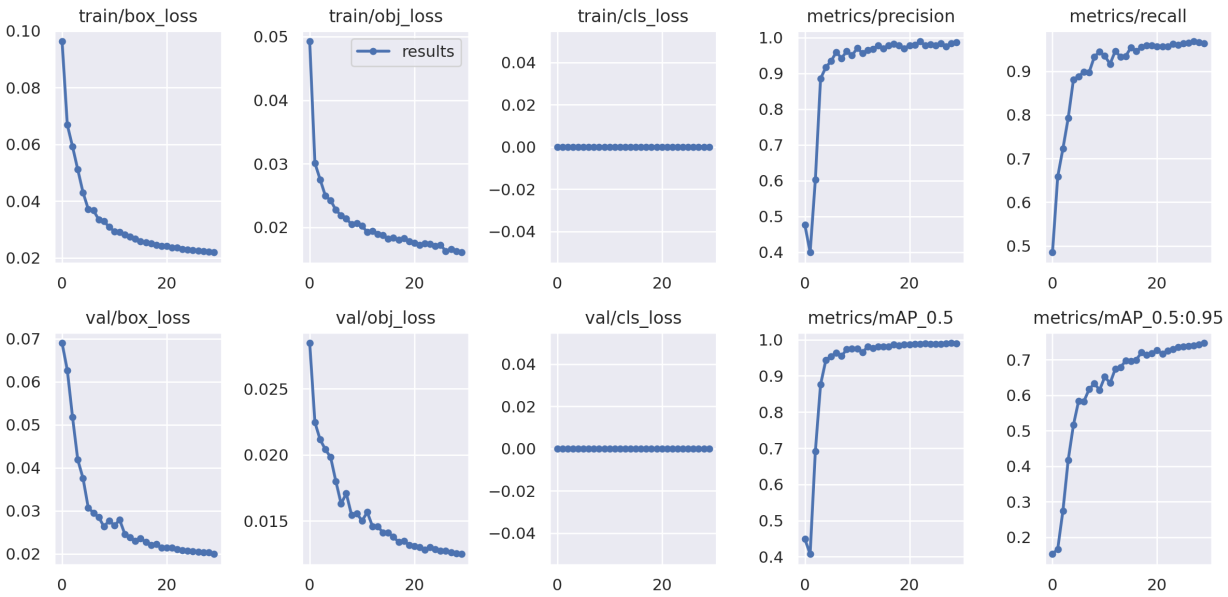


2. Individual training curves for different network dimensions.

3. Worm Detection Dataset: Consists of 1407 images for training, 180 for validation and 171 for testing Deep-Worm-Tracker.

4. Worm re-ID Dataset: the dataset consists of 32 worm identities and is structured according to the MARS re-ID dataset.

5. Worm re-ID models checkpoint files

6. Worm Test files used for testing Deep-Worm-Tracker. Worm video recordings were captured using the image acquisition setup mentioned earlier.

7. Tracking results of the Deep-Worm-Tracker.

8. Google Colab notebook for a quick start for running Deep-Worm-Tracker after cloning from github.
